# A Statistical Framework for Data Purification with Application to Microbiome Data Analysis

**DOI:** 10.1101/2021.09.13.460157

**Authors:** Zequn Sun, Jing Zhao, Zhaoqian Liu, Qin Ma, Dongjun Chung

## Abstract

Identification of disease-associated microbial species is of great biological and clinical interest. However, this investigation still remains challenges due to heterogeneity in microbial composition between individuals, data quality issues, and complex relationships among species. In this paper, we propose a novel data purification algorithm that allows elimination of noise observations, which leads to increased statistical power to detect disease-associated microbial species. We illustrate the proposed algorithm using the metagenomic data generated from colorectal cancer patients.

## Introduction

Trillions of microbes, including bacteria, archaea, viruses, and eukaryotic microbes, reside in and on human bodies. They involve various crucial physiological processes of human, such as influencing immunity and regulating metabolism [1]. Back in the 19th century, a seminal study has demonstrated that shifts in microbial composition were tightly associated with diseases [2]. Since then, increasing effort has been made to explore disease-relevant microbes [2], thereby contributing to clinical diagnostic and treatment of diseases. However, the research was seriously hampered since only less than 1% of microbes can be studied based on traditional independent-culture methods [3].

The significant advance in sequencing technologies paves the way for a comprehensive understanding of microbial contributions to diseases, such as 16S ribosomal RNA sequencing and whole-genome shotgun metagenomic sequencing [3]. In contrast to independent-culture technologies, these methods enable insights into large amounts of non-cultivable microbes [3]. Currently, multiple nationwide projects have been established, such as the Human Microbiome Project [4], the Interactive Human Microbiome Project [5], and the Metagenomics of the Human Intestinal Tract project [6], which generated a massive amount of microbial data from tissue swabs or stool samples of health cohorts and diseases (e.g., prediabetes and inflammatory bowel diseases). Recently, Thomas et al. constructed an integrated metagenomic dataset of the human gut microbiome for colorectal cancer (CRC) studies [7]. Additionally, Dohlman et al. re-examined whole-genome and whole-exon sequencing studies in The Cancer Genome Atlas and collected microbial reads for oropharyngeal, esophageal, gastrointestinal, and colorectal tissues [8], providing a distinct view on elucidating microbial composition of internal organs to various cancers. Together, these available data offer an unprecedented opportunity to reveal disease-associated microbial species (a.k.a., microbial signatures).

However, identifying reliable associations between human microbial signatures with diseases is still a challenging task [9]. The heterogeneity in microbial composition between individuals leads to an increased probability of spurious associations due to the confounding effects of human variables [10]. Geographical location as one of the human variables has been shown to influence the linkage between microbial composition and an increased risk of metabolic disease [11]. Meanwhile, multiple studies using 16s rRNA gene or metagenomic sequencing had linked discordant microbial species with CRC individually, and the evaluation of the generalizability of these microbial signatures in an expanded population is emerging [7, 12–17]. Meta-analyses performed on multiple large-scale cohorts based on metagenomic data have demonstrated that the generalizability of cross-study can be improved by including additional healthy samples, while the models transferred between studies were less accurate than the performance within-studies [18, 19]. Later on, robust global microbiome signatures were identified and validated across multiple cohorts, where the predictive and functional roles of these signatures were examined [7, 20, 21].

From the statistical point of view, this study can be considered as a data purification problem. Specifically, we assume that our data consists of signal and noise observations and we aim to improve statistical power to detect disease-associated microbial species by either eliminating noise observations or over-weighting signal observations (equivalently, down-weighting noise observations). However, this analysis is not a trivial task because it is hard to know signal and noise observations *a priori* and there are complex inter-relationships among microbial species. In this paper, we propose a novel algorithm to improve microbial species detection through data purification based on a nonparametric approach and a subsampling technique. We illustrate the proposed algorithm and its performance using metagenomic data obtained from CRC patients.

## Methods

**Algorithm 1** describes the proposed algorithm. We denote the observation index as *i,i* = 1,⋯, *n*, and the species index as *j, j* = 1,⋯, *p*. First, Step 1 provides the baseline statistical power for species identification when no data purification is implemented. Using this baseline, we can evaluate gain or loss of including/excluding each observation in the data in the following steps. Second, in Step 2, we evaluate the quality of each observation in a nonparametric way. Specifically, we sample *k* observations without replacement from the data multiple times and identify species using each subsampled data. Because each subsampling includes and excludes observations in different ways, this approach allows us to consider various scenarios with slightly different modification of the data. For *i*-th observation in *j*-th subsample, we have four possibilities: (*i*) an observation is included (*T_ij_* = 1) and more species are identified (*X_j_* = +1); (*ii*) an observation is included (*T_ij_* = 1) and less species are identified (*X_j_* = −1); (*iii*) an observation is not included (*T_ij_* = 0) and more species are identified (*X_j_* = +1); and (*iv*) an observation is not included (*T_ij_* = 0) and less species are identified (*X_j_* = −1). Note that (i) and (iv) indicate that statistical power to detect species can potentially be increased by including this observation. In contrast, (ii) and (iii) indicate that statistical power to detect species can be rather decreased by including this observation.

Based on this rationale, in Step 3, we construct the important score for *i*-th observation as *S_i_* = {*Y_i++_ + Y_i−−_ – Y_i+−_ – Y_i−{_*}/*m*, where *m* is the number of subsampling, 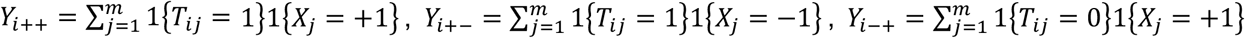, and 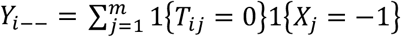. Again, note that this important score increases as there are more subsamples with increased statistical power by including this observation (case i), or with decreased statistical power by not having this observation (case iv). Likewise, this important score also increases as there are less subsamples with decreased statistical power by including this observation (case ii), or with increased statistical power by not having this observation (case iii). Finally, in Step 4, we order observations based on their importance scores, choose the top *d* observations, and use them for the final species detection.

In this algorithm, there are three tuning parameters, subsample size (*k*), number of subsampling (*m*), and the sample size used for final species detection (*d*). The first two parameters affect accuracy of estimating usefulness of each observation and accuracy of calculating the importance score, respectively. Based on the rationale that bootstrapping is close to 63.2% sampling without replace, we set *k* close to 0.632*n*. In the case of number of subsampling, it suffices to make it large enough and we use 10,000 by considering both computing accuracy and efficiency. The sample size used for final species detection (*d*) affects the performance more significantly as it directly controls the degree of data purification. We investigate this parameter more in depth in the Results section.

### Algorithm 1

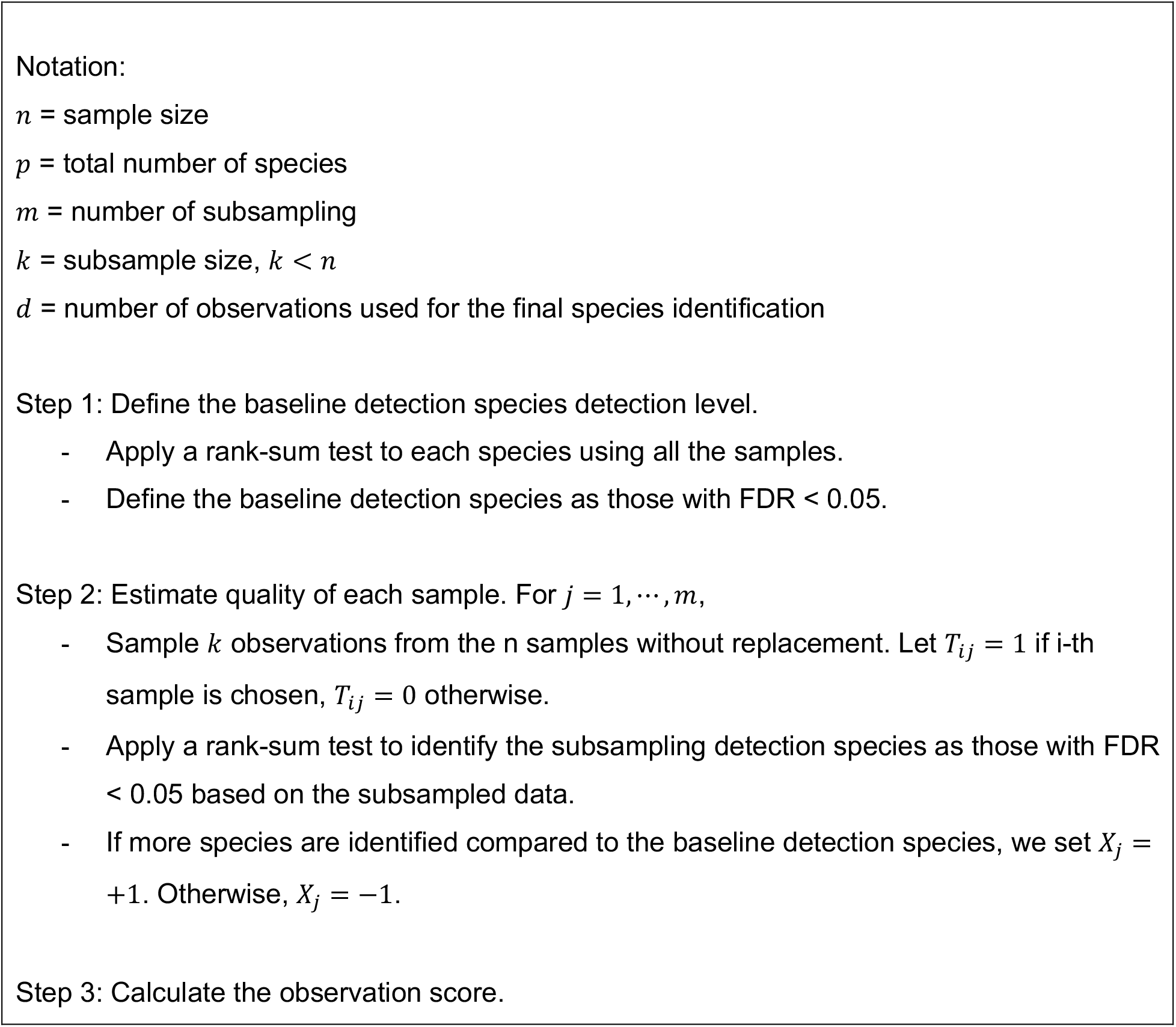
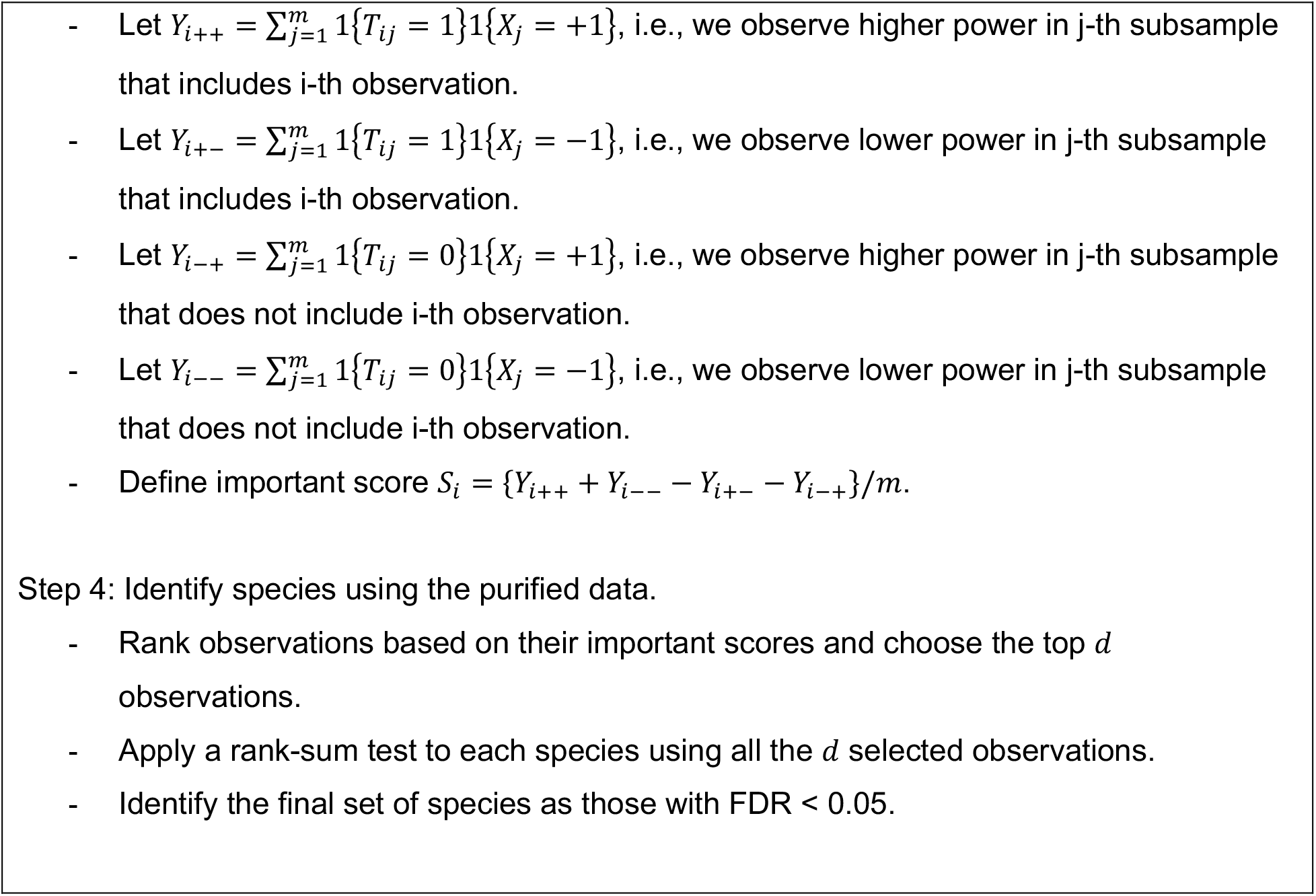

## Results

For the real data analysis of microbiome data, we downloaded the preprocessed metagenomic data of six cross-sectional studies of CRC from the curatedMetagenomicData R package [22], which is supplemental to the meta-analysis by Thomas, A. M., et al. [7]. The data includes information of metadata, taxonomic and functional composition, and relative abundance of each species and metabolic pathway. Since the preprocessed metagenomic data for the other two validation studies used in the meta-analysis were not available, we downloaded their raw fastq files from the DNA Data Bank of Japan database (project No. DRA00668432) and European Nucleotide Archive (project No. PRJEB2792814), respectively. Data preprocessing was performed strictly as described in the meta-analysis to make it consistent with the other six cohorts.

Overall, we use the name of the country where the cohorts were recruited to denote the eight studies: ThomasAM_2018a represents cohort in Italy (ITA 1), ThomasAM_2018b represents cohort in Italy (ITA 2), FengQ_2015 represents cohort in Austria (AUS), VogtmannE_2016 represents cohort in the United States (USA), YuJ_2015 represents cohort in China (CHI), and ZellerG_2014 represents cohort in France (FRA), project No. DRA006684 represents cohort in Japan (JAP), and project No. PRJEB27928 represent cohorts in Germany (GEM). In total, the pooled samples contain 387 CRC cases and 384 healthy controls, while there are the total 695 species in this dataset.

First, we investigate the impact of tuning parameters on the performance in the sense of identified species. **Figure 1** shows the number of identified species as a function of the sample size (*d*) and the subsampling size (*m*). We can observe similar curve shapes across different subsampling sizes (*m*) but overall, the curves move up until we use subsampling sizes of 400 to 500. Note that here 0.632*n* = 487.3, which is located between 400 and 500. This supports our choice of setting *m* = 0.632*n*. Within each panel, we can see that we have the highest statistical power when we use the sample size (*d*) of 300 or 400, which is approximately the half of the total number of species (*p*). Based on this result, we focus on results based on the top half observations (*d* = 0.5*n*).

**Figure 1.**
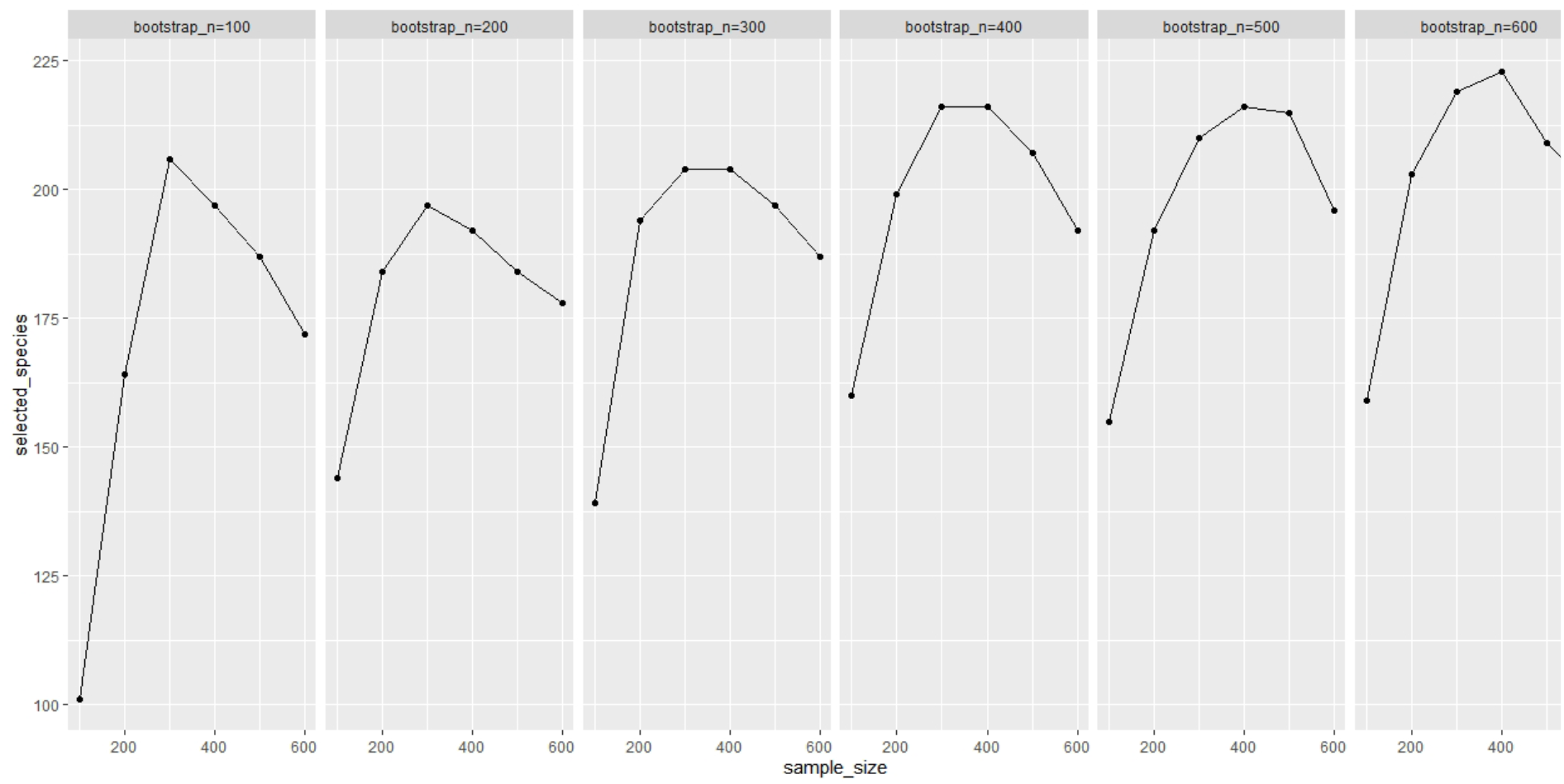
The number of identified species as a function of the sample size (d; x-axis) and the subsampling size (m; panel).

Second, to evaluate the performance of selecting “good” observations (i.e., data purification), we applied the proposed algorithm, obtained the observation subset based on importance scores, and checked from which cohort more observations are selected. To evaluate the quality of each cohort, we applied a rank-sum test to each cohort separately and checked how many species were identified using each cohort. Based on this criterion, we found that, among the 8 cohorts we considered in this study, PRJDB4176 (1% species identified), ThomasAM_2018a (0% species identified), ThomasAM_2018b (1% species identified), and VogtmannE_2016 (0% species identified) are noisier compared to YuJ_2015 (5% species identified), ZellerG_2014 (1% species identified), FengQ_2015 (6% species identified), and PRJEB27928 (17% species identified). **Table 1** shows the number of observations selected from each cohort for different *d* parameter. We can see that more observations are selected from less noisy cohorts (the last four rows). Moreover, the fold change of selected observations between the less noisy cohorts (the last four rows) vs. the noisier cohorts (the first four rows) is largest when *d* is 300 or 400, which again supports our choice of this parameter.

**Table 1.** Number of observations selected from each cohort for different d parameter.

| Cohorts | Sample size | d=100 | d=200 | d=300 | d=400 | d=500 | d=600 |
| --- | --- | --- | --- | --- | --- | --- | --- |
| PRJDB4176 | 80 | 7 | 13 | 29 | 44 | 54 | 62 |
| ThomasAM_2018a | 53 | 3 | 8 | 14 | 22 | 34 | 43 |
| ThomasAM_2018b | 60 | 9 | 17 | 19 | 26 | 32 | 38 |
| VogtmannE_2016 | 104 | 4 | 13 | 25 | 41 | 59 | 77 |
| YuJ_2015 | 128 | 20 | 39 | 50 | 75 | 90 | 106 |
| ZellerG_2014 | 114 | 17 | 29 | 49 | 58 | 71 | 85 |
| FengQ_2015 | 107 | 25 | 41 | 54 | 61 | 70 | 81 |
| PRJEB27928 | 125 | 16 | 44 | 63 | 76 | 94 | 108 |

Finally, we investigated the 226 species associated with CRC that were identified by the proposed algorithm. **Figure 2** shows the phylogenetic relationships of the species. They are dominated by *Firmicutes, Bacteroidetes, Proteobacteria, Actinobacteria,* and *Fusobacteria,* which is consistent with previous studies about CRC-associated microbial signatures [23, 24]. We further collected 13 reliable CRC-associated species from two up-to-date meta-analysis studies of CRC (those have been demonstrated associated with CRC by both the two studies) [7, 21], and observed all these species have been detected by our method, demonstrating the high effectiveness of our method. Notably, the proposed method found 104 novel species compared to the baseline species list, including phyla that have not been studied with CRC before, such as *Candidatus Saccharibacteria* with two species (candidate division TM7 single-cell isolate TM7b and candidate division TM7 single-cell isolate TM7c). These can provide insights into CRC signature studies.

**Figure 2.**
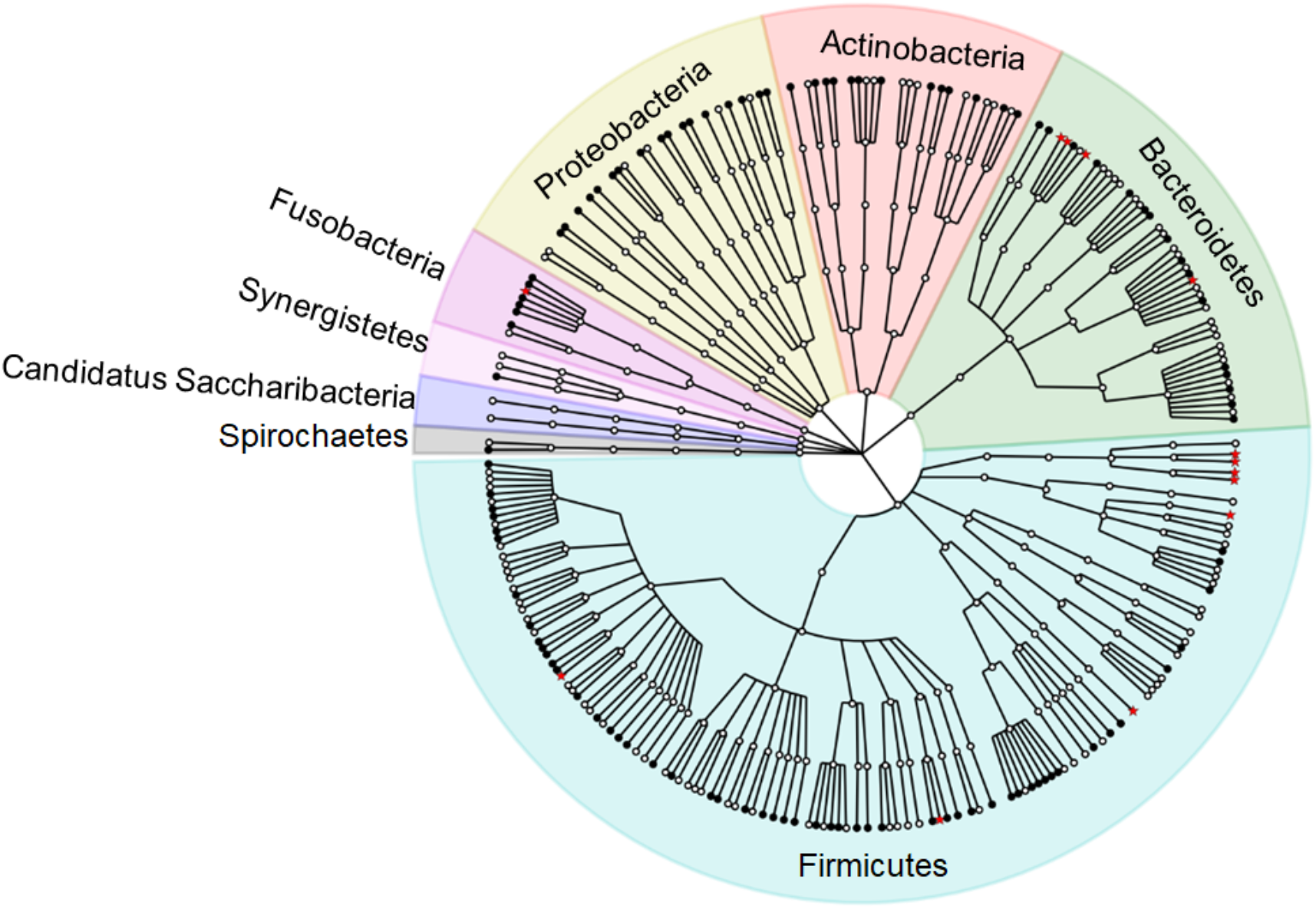
The phylogenetic tree of 226 species identified from the proposed algorithm. From inner to outer, each ring consisting of multiple nodes represents a taxonomic level, from phylum to species, respectively. Specifically, each node on the outmost ring represents a species. Different colors indicate different phylum. The nodes as black dots represent the overlapped species with baseline list species (totally 122 here), while the red stars represent the 13 species associated with CRC from the previous two studies.

## Discussion

In this paper, we propose a novel algorithm to improve detection of disease-associated microbial species by purifying the data (i.e., remove noise observations). The proposed algorithm is based on a nonparametric approach and a subsampling technique. This helps address complex data structure we often observe in microbiome data. Our application to the CRC metagenomics data indicates that the proposed algorithm can potentially lead to improved statistical power to identify key microbial species. However, the proposed algorithm also has limitations as well. Our current study focuses on the disease-related microbial signatures by comparing different cohort populations. This may lead to spurious associations with diseases since different taxonomic compositions of microbes may contribute to similar functionality [25]. With the available metagenome and metatranscriptome data [26, 27], one can give a read-out of functional activities within microbial communities, not limited to taxa, thereby studying the disease-associated taxonomic and functional mechanisms simultaneously. Notably, the high sparsity, noise, and heterogeneity are intrinsic characteristics of microbial data [28]. One promising direction in future research is introducing deep learning models, which have shown strong power in learning knowledge from large-scale sparse and noisy data [29], and we currently investigate this direction to further improve detection of disease-associated microbial species.

## References

1. Sepich-Poore, G.D., et al., The microbiome and human cancer. Science, 2021.371(6536): p. eabc4552.

2. Dethlefsen, L., M. McFall-Ngai, and D.A. Relman, An ecological and evolutionary perspective on human–microbe mutualism and disease. Nature, 2007. 449(7164): p. 811–818.

3. Riesenfeld, C.S., P.D. Schloss, and J. Handelsman, Metagenomics: genomic analysis of microbial communities. Annual review of genetics, 2004. 38: p. 525–552.

4. Methé, B.A., et al., A framework for human microbiome research. Nature, 2012. 486(7402): p. 215–221.

5. Proctor, L.M., et al., The Integrative Human Microbiome Project. Nature, 2019. 569(7758): p. 641–648.

6. Qin, J., et al., A human gut microbial gene catalogue established by metagenomic sequencing. Nature, 2010. 464(7285): p. 59–65.

7. Thomas, A.M., et al., Metagenomic analysis of colorectal cancer datasets identifies cross-cohort microbial diagnostic signatures and a link with choline degradation. Nature Medicine, 2019. 25(4): p. 667–678.

8. Dohlman, A.B., et al., The cancer microbiome atlas: a pan-cancer comparative analysis to distinguish tissue-resident microbiota from contaminants. Cell Host Microbe, 2021. 29(2): p. 281–298.e5.

9. Elinav, E., et al., The cancer microbiome. Nature Reviews Cancer, 2019. 19(7): p. 371–376.

10. Vujkovic-Cvijin, I., et al., Host variables confound gut microbiota studies of human disease. Nature, 2020. 587(7834): p. 448–454.

11. He, Y., et al., Regional variation limits applications of healthy gut microbiome reference ranges and disease models. Nature Medicine, 2018. 24(10): p. 1532–1535.

12. Kostic, A.D., et al., Fusobacterium nucleatum potentiates intestinal tumorigenesis and modulates the tumor-immune microenvironment. Cell Host Microbe, 2013. 14(2): p. 207–15.

13. Rubinstein, M.R., et al., Fusobacterium nucleatum promotes colorectal carcinogenesis by modulating E-cadherin/β-catenin signaling via its FadA adhesin. Cell Host Microbe, 2013. 14(2): p. 195–206.

14. Yu, J., et al., Metagenomic analysis of faecal microbiome as a tool towards targeted non-invasive biomarkers for colorectal cancer. Gut, 2017. 66(1): p. 70–78.

15. Feng, Q., et al., Gut microbiome development along the colorectal adenoma–carcinoma sequence. Nature Communications, 2015. 6(1): p. 6528.

16. Zeller, G., et al., Potential of fecal microbiota for early-stage detection of colorectal cancer. Molecular Systems Biology, 2014. 10(11): p. 766.

17. Kostic, A.D., et al., Genomic analysis identifies association of Fusobacterium with colorectal carcinoma. Genome Res, 2012. 22(2): p. 292–8.

18. Pasolli, E., et al., Machine Learning Meta-analysis of Large Metagenomic Datasets: Tools and Biological Insights. PLOS Computational Biology, 2016. 12(7): p. e1004977.

19. Dai, Z., et al., Multi-cohort analysis of colorectal cancer metagenome identified altered bacteria across populations and universal bacterial markers. Microbiome, 2018. 6(1): p. 70.

20. Marcos-Zambrano, L.J., et al., Applications of Machine Learning in Human Microbiome Studies: A Review on Feature Selection, Biomarker Identification, Disease Prediction and Treatment. Frontiers in Microbiology, 2021. 12(313).

21. Wirbel, J., et al., Meta-analysis of fecal metagenomes reveals global microbial signatures that are specific for colorectal cancer. Nature Medicine, 2019. 25(4): p. 679–689.

22. Pasolli, E., et al., Accessible, curated metagenomic data through ExperimentHub. Nature Methods, 2017. 14(11): p. 1023–1024.

23. Pan, H.W., et al., Biodiversity and richness shifts of mucosa-associated gut microbiota with progression of colorectal cancer. Res Microbiol, 2020. 171(3-4): p. 107–114.

24. Sinha, R., et al., Fecal microbiota, fecal metabolome, and colorectal cancer interrelations. PloS one, 2016. 11(3): p. e0152126.

25. Snipen, L., et al., Reduced metagenome sequencing for strain-resolution taxonomic profiles. Microbiome, 2021.9(1): p. 79.

26. Lloyd-Price, J., et al., Multi-omics of the gut microbial ecosystem in inflammatory bowel diseases. Nature, 2019. 569(7758): p. 655–662.

27. Abu-Ali, G.S., et al., Metatranscriptome of human faecal microbial communities in a cohort of adult men. Nature Microbiology, 2018. 3(3): p. 356–366.

28. Zhou, F., et al., Bayesian biclustering for microbial metagenomic sequencing data via multinomial matrix factorization. arXiv preprint arXiv:2005.08361,2020.

29. Hu, Z., et al., Heterogeneous Graph Transformer, in Proceedings of The Web Conference 2020. 2020, Association for Computing Machinery: Taipei, Taiwan. p. 2704–2710.

